# Beyond the surface: plasmalogens are dispensable for retinal integrity and fertility in the mouse

**DOI:** 10.64898/2026.04.09.717382

**Authors:** Ilaria Dorigatti, Viktorija Juric, Michael J. Blumer, Denise Kummer, Janik Kokot, Georg Golderer, Fabian Dorninger, Johannes Berger, Markus A. Keller, Katrin Watschinger

**Affiliations:** Institute of Molecular Biochemistry, Biocenter, Medical University of Innsbruck, 6020 Innsbruck, Austria; Institute of Human Genetics, Medical University of Innsbruck, 6020 Innsbruck, Austria; Department of Anatomy, Histology and Embryology, Institute of Clinical and Functional Anatomy, Medical University of Innsbruck, 6020 Innsbruck, Austria; Department of Pathobiology of the Nervous System, Center for Brain Research, Medical University of Vienna, 1090 Vienna, Austria

**Keywords:** ether lipids, PEDS1, Rhizomelic Chondrodysplasia Punctata, lipidomics, cataract

## Abstract

Ether lipids and their subclass, the plasmalogens, are critical regulators of membrane organization, signaling, and stress responses in multiple tissues. Inborn errors in their anabolism cause severe multi-organ diseases such as Rhizomelic Chondrodysplasia Punctata and related peroxisomal disorders. The *Gnpat* knockout mouse model, characterized by total ether lipid deficiency, recapitulates key features of this disorder, including dense bilateral cataracts, microphthalmia, and infertility, but the specific contribution of different subclasses like plasmalogens remains elusive. The recent identification of the *Peds1* gene allows dissecting the impact of selective plasmalogen deficiency with retention of plasmanyl lipids, another ether lipid subgroup. Here, we performed the first side-by-side comparison of *Gnpat* and *Peds1* knockout in mice on a matched genetic background (C57BL/6 x CD1). In contrast to the situation in *Gnpat* knockout mice, plasmanyl lipids in *Peds1* knockout mice were sufficient to prevent cataract formation and maintain normal ocular structures, despite marked shifts in the ocular phospholipidome. Also, fertility and reproductive function were found to be preserved in *Peds1* knockout mice. Our data demonstrate that plasmanyl lipids can partially protect against the severe phenotypes observed in mouse models of total ether lipid deficiency; notably, the ocular and reproductive phenotypes were plasmalogen-independent, indicating that loss of the vinyl ether double bond is not the key determinant of all symptoms in human and murine ether lipid deficiency and can at least partly be compensated by plasmanyl lipids.

**Highlights:**

- In mice, total ether lipid deficiency causes cataracts and infertility.
- The role of plasmalogens in these phenomena remains unclear.
- Two *PEDS*1-deficient patients were reported, but cataracts were observed in only one case.
- *Peds*1-deficient mice have no cataracts or ocular abnormalities.
- Mice with a deletion of *Peds1* display normal fertility rates.

## 1. Introduction

Ether lipids including the subclass of plasmalogens have emerged as key regulators of membrane organization, signaling, and stress responses in diverse tissues [1]. Their clinical relevance is underscored by inborn errors of ether lipid biosynthesis that cause multi-organ disease, specifically Rhizomelic Chondrodysplasia Punctata (RCDP) and related peroxisomal disorders [1,2]. Although associations with neurodegeneration, including Alzheimer’s disease, have been proposed, causal relations remain incompletely defined [3].

The ether lipid biosynthesis pathway is segregated into two organelles. In peroxisomes, the pathway initiates with the action of glyceronephosphate *O*-acyltransferase (GNPAT; EC 2.3.1.42), followed by alkylglycerone phosphate synthase (AGPS; EC 2.5.1.26), with fatty acyl-CoA reductases FAR1/FAR2 (EC 1.2.1.84) supplying the required fatty alcohols. Subsequent maturation occurs at the cytosolic leaflet of the endoplasmic reticulum (ER), where plasmanylethanolamine desaturase (PEDS1; EC 1.14.19.77) introduces the defining vinyl ether double bond, converting plasmanyl (alkyl) phospholipids to plasmenyl (vinyl ether) lipids (better known as plasmalogens) [4].

Two subtypes of RCDP arise from mutations in peroxins (PEX): i) type 1 (OMIM 601757) caused by mutations in *PEX7*, which is needed for the correct import of AGPS and other PTS2 cargos into peroxisomes, and ii) type 5 (OMIM 600414 and 616716), which is evoked by the deficiency in PEX5L, a co-receptor for PEX7. Pathogenic variants in *GNPAT* (RCDP type 2, OMIM 602744 and 222765) and *AGPS* (RCDP type 3, OMIM 603051 and 600121) lead to further subtypes of this disease, while mutations in *FAR1*, although associated with a similar pattern of neurodevelopmental delay and cataracts, lack the skeletal phenotype and are therefore classified as peroxisomal fatty acyl-CoA reductase 1 disorder (PFCRD, OMIM 616154).

Across RCDP types 1, 2, 3 and 5 and PFCRD, early-onset bilateral congenital cataracts and additional ocular anomalies are frequently reported, alongside facial features such as hypertelorism and epicanthal folds [5,6]. Notably, in the first and very recently described individuals with *PEDS1* deficiency, congenital cataracts were observed in only one of the two cases [7], suggesting lower penetrance upon selective plasmalogen deficiency compared with total ether lipid deficiency.

The eye is particularly rich in ether lipids. Plasmalogens and plasmanyl lipids are abundant in retinal membranes and optic nerve myelin [8], where region-specific enrichment patterns have been described, including ethanolamine plasmalogens (PE(P)) species with monounsaturated fatty acids in the optic nerve and DHA-containing PE(P-18:0/22:6) in the inner retina [9]. Ether lipids were found to be synthesized in the inner segment of photoreceptor cells and in the retinal pigment epithelium (RPE) [10] with PE(P) (major species: PE(P-16:0/20:4), PE(P-18:0/20:4) and PE(P-18:0/22:6)) being present in the retina and RPE cells and choline plasmalogens (PC(P)) (major species: PC(P-16:0/18:0), PC(P-18:0/18:0) and PC(P-18:1/18:0)) in the optic nerve. Their levels were described to decline with age [11]. Also, lens epithelial cells have the ability to synthesize ether-linked lipids [12] and significant amounts of these lipids were found in murine [12] and human [13] lenses, where approximately one-third of phospholipids are ether-linked, with a predominance of ethanolamine- and serine-linked plasmanyl lipids; plasmanylethanolamine PE(O-18:1/18:1) has been reported as a highly abundant molecular species [13]. The plasmalogen PE(P-18:1/18:1) was found to be the second most abundant glycerophospholipid in mouse lenses [14]. Plasmanyl ether lipids are reduced in cataractous lenses [13]. Together, these observations implicate a role of ether lipids in ocular structure and function.

Historically, dissection of vinyl ether-dependent biology has been limited by the unknown identity of PEDS1 and by analytical challenges in distinguishing (P) from (O) species. The identification of *TMEM189* as gene coding for PEDS1 and advances in liquid chromatography-mass spectrometry (LC-MS) now enable reliable resolution separating plasmalogens from plasmanyl lipids [15–18]. Recent evidence from cell and tissue systems suggests that some functions attributed to plasmalogens can be partially taken over by plasmanyl counterparts [see preprint 19], whereas others uniquely depend on the vinyl ether double bond (e.g., antioxidant reactivity and specific biophysical properties) [4,20], potentially explaining phenotypic variability across defects evoked by ether lipid deficiency. Importantly, two canonical plasmanyl lipids, platelet-activating factor and seminolipid, underscore that non-plasmalogen ether lipid species can have nonredundant physiological roles independent of plasmalogens [21].

Mouse models recapitulate key aspects of human disease and offer a tractable platform to separate total ether lipid deficiency from isolated plasmalogen loss. Knockouts (KO) of peroxisomal enzymes in the mouse (e.g., *Gnpat* KO, *Agps* KO, [22,23]) lack essentially all ether lipids and, among a variety of other abnormalities, develop early cataracts and additional ocular phenotypes like microphthalmia, including retinal vascular anomalies and “glaucoma-like” optic nerve changes [8,23–25]. Male infertility is a consistent feature across totally ether lipid-deficient mouse lines, was attributed to the loss of plasmalogens and consequent defects in spermatogenesis and sperm maturation; female reproductive effects are milder [23–27] but in a recent study *Gnpat* KO females have been reported to be unable to reproduce [28]. Data on the eye phenotypes of *Far1*-deficient mice remain sparse despite recent model descriptions [26,29], and a Pex5L-specific KO model with RCDP-like features has not been reported to date. By contrast, *Peds1* KO mice, which selectively lack plasmalogens but retain plasmanyl lipids, have recently become available [16]; initial large-scale phenotyping suggests partial preservation of fertility and an abnormal eye and retina morphology and pigmentation, but detailed ocular and reproductive assessments are limited (https://www.mousephenotype.org/data/genes/MGI:2142624).

Here, we address this gap by directly comparing two complementary mouse models on a matched genetic background (C57BL/6 × CD1): *Gnpat* KO (total ether lipid deficiency) and *Peds1* KO (selective plasmalogen deficiency, with retention of plasmanyl lipids). We test the hypothesis that plasmanyl lipids can preserve key ocular and reproductive functions in the absence of plasmalogens. To this end, we combine targeted lipidomics capable of distinguishing PE(P) from PE(O) species with assessments of fertility and eye structure. This design enables a systematic evaluation of which phenotypes can be assigned to the lack of the vinyl ether double bond and which can be buffered by the presence of plasmanyl ether lipids.

## 2. Material and methods

### 2.1. Ethics statement and animal models

All animal breedings were approved by the Austrian Federal Ministry of Education, Science and Research (BMBWF-66.011/0100-V/3b/2019, 2024-0.307.678, 2023-0.088.476, 2021-0.556.156 and 2024-0.802-540).

*Gnpat* KO mice (*Gnpat^tm1Just^*; MGI:2670462) have been described previously [24] and were maintained on an outbred C57BL/6 × CD1 background. The *Peds1* KO mouse line (strain Tmem189tm1a(KOMP)Wtsi) on C57BL/6N background was obtained from the Wellcome Sanger Institute (Hinxton, Cambridge, UK) and distributed via EMMA/Infrafrontier GmbH (Munich, Germany) [30–33]. To enable direct comparison of the phenotypes between the two models, *Peds1* KO mice were backcrossed to the mixed C57BL/6 x CD1 genetic background of *Gnpat* KO mice [24]. Genotyping was performed as described previously [24,34]. Breeding was routinely conducted by heterozygous mating; a limited number of KO x KO pairings were set up to assess fertility in the *Peds1* KO line only. Mice were kept in specific pathogen free (SPF) conditions in individually ventilated cages with nesting material, under a controlled 12 hours light/12 hours dark cycle at 21°C and 50% humidity, with *ad libitum* access to standard chow and water. Animals were sacrificed by cervical dislocation and tissues were either snap-frozen in liquid nitrogen and stored at -80°C until biochemical and lipidomic analyses, or were collected and fixed in 4% paraformaldehyde (PFA) for histological analysis. The genotypes of experimental animals were confirmed *post mortem*.

### 2.2. PEDS1 enzymatic activity in whole eyes

Microsomal fractions were prepared by homogenizing the eyes in 200 µl 0.1 mM Tris HCl and 0.25 M sucrose (pH 7.2), with a cocktail of protease inhibitors (aprotinin 50 µg/ml; pepstatin A 50 µM; trypsin inhibitor 5 mg/ml; iodoacetamide 100 mM; leupeptin 50 µg/ml), using an Ultra Turrax (IKA, Staufen, Germany) followed by centrifugation at 3,000 x g for 10 min at 4°C. The supernatant was centrifuged at 20,000 x g for 30 min at 4°C, and the pellet was resuspended in 30 µl Tris-HCl-sucrose buffer, including protease inhibitors, and protein concentration was quantified by Bradford assay (Bio-Rad, Hercules, California, USA, #5000006), using BSA as standard. The suspension was adjusted with 3.5 mg/ml BSA in PBS including protease inhibitors, to a final concentration of 0.5 mg microsomal protein per ml.

PEDS1 activity assay was performed as previously described using purified 1-*O*-pyrenedecyl-*sn*-glycero-3-phosphoethanolamine as substrate [35]. Activity was calculated from the HCl-liberated pyrenedecanal peak with fluorescence detection at 340/405 nm; separations were run on a Zorbax Eclipse XDB-C8 column using the published gradient on an 1200 HPLC system (Agilent Technologies, Vienna, Austria) equipped with a thermostatted autosampler, UV-Vis, fluorescence detectors and a column thermostat.

### 2.3. Quantification of plasmalogen content in whole eyes

Frozen eyes were homogenized in 200 µl PBS with an Ultra-Turrax (IKA, Staufen, Germany). Protein concentration was determined by Bradford assay (see above), and all homogenates were adjusted to a concentration of 0.5 mg/ml. Aqueous homogenate (200 µl) was extracted twice with 500 μl chloroform/methanol (2:1 v/v). Combined organic phases were evaporated to dryness and reconstituted in 100 μl acetonitrile (ACN)/ethanol (1:1 v/v) with constant agitation for 10 min at 37°C. Plasmalogens were cleaved and aldehydes released by incubation of 10 µl of lipid extract with 40 μl dansylhydrazine (0.45 mg/ml, Merck, 03334) in ACN/2 M HCl (930:70 v/v). In parallel, for quantification of free aldehydes derived from other cellular sources, 10 µl extract was incubated with 40 μl dansylhydrazine (0.45 mg/ml) in ACN/2 M acetic acid (930:70 v/v). After 15 min on ice in the dark, both samples were centrifuged for 5 min at 20,000 x g, and 10 µl of each supernatant was injected onto the same Agilent HPLC system under the same running conditions as described in chapter 2.2.. Dansylhydrazones eluted at 8-10 min were quantified by fluorescence peak area (excitation 340 nm, emission 525 nm). Final plasmalogen levels were calculated by correcting the total aldehyde signal for the minor contribution of free aldehydes (typically less than 1%). This method follows a published protocol [35].

### 2.4. Lipid extraction and mass spectrometrical analysis of the murine ocular phospholipidome

Homogenization and protein quantification was performed as in chapter 2.3., and all homogenates were adjusted to a concentration of 1 mg/ml. 200 µl aqueous homogenate was extracted twice with 500 μl chloroform/methanol (2:1 v/v) containing 2.5 µl SPLASH™ LIPIDOMIX™ Mass Spec Standard 330707 internal standard mix (Avanti Polar Lipids, Albaster, AL, USA). Combined organic phases were evaporated to dryness under a stream of nitrogen and overlaid with argon to prevent oxidation.

Extracted lipids were reconstituted in 100 µl of isopropanol/acetonitrile/water (IPA/ACN/H₂O, 54:30:16, v/v/v) and analyzed on a timsTOF Pro times-of-flight mass spectrometer (Bruker Daltonics, Bremen, Germany) coupled to an Elute HPLC system (Bruker Daltonics). Ionization was achieved in negative mode with HESI (Vacuum Insulated Probe Heated Electrospray Ionization). InfinityLab Poroshell 120 EC-C18, 2.1 × 100 mm, 2.7 µm was used for chromatographic separation of the lipids. The column was maintained at 55°C with the flow-rate of 0.6 ml/min as described previously [36], using the mobile phase composition and gradient according to [37]. Data were acquired in parallel accumulation-serial fragmentation mode with data-dependent MS/MS acquisition (DDA) and ion mobility separation. Key acquisition parameters were: MS1 and MS2 mass ranges m/z 50-1700. Ion mobility range was set from 0.55 to 1.9 V·s/cm² with IMS Ramp Time 100.0 ms. Source parameters were: capillary 4500 V, nebulizer pressure 2.5 bar, dry gas 8.0 l/min, dry heater 260°C, multipole RF 200.0 Vpp and collision RF 1100.0 Vpp. The instrument was calibrated with a solution of sodium formate in a High Precision Calibration (HPC) calibration method. Raw data files were processed by using MS-DIAL (v. 4.92) [38]. LipidBlast *in a silico-generated* database was used for accurate mass and MS/MS matching [39]. MS/MS spectra for all detected features were checked and confirmed manually, resulting in 160 high-confidence phospholipid features. Lipid class-specific elution patterns were systematically inspected based on their retention time and m/z values. Additionally, retention behavior for plasmalogens and plasmanyl species were annotated according to the reference library described in [40]. Processed data with confirmed species was exported from the MS-DIAL. MetaboAnalyst 6.0. was used to normalize the data to sum, and to generate heatmap and principal component analysis (PCA) plots [41].

### 2.5. Histological analysis of murine eyes

The eyes, including the optic nerve, of male and female mice (wildtype (WT) (on C57BL/6 x CD1 and C57BL/6N background), as well as *Peds1* KO and *Gnpat* KO (on C57BL/6 x CD1 background) aged 5 weeks and 3 months were fixed with 4% PFA in 0.1 M phosphate-buffered saline (PBS, pH = 7.4) at 4°C for 12 hours. The samples were then rinsed in PBS, dehydrated in a graded series of isopropanol and xylene, and embedded in paraffin using a routine histological infiltration processor (Miles Scientific Inc., Naperville, IL, USA). Sagittal serial sections (7 µm) were cut using an HM 355 S microtome (Microm, Walldorf, Germany) and mounted on glass slides. The sections were deparaffinized, stained with hematoxylin (8 min) and eosin (2 min) (H&E), dehydrated, and coverslipped. They were examined with a Zeiss ax10 microscope equipped with a Zeiss AxioCam 512 color digital camera and ZEN 3.0 blue edition software (Zeiss, Oberkochen, Germany).

### 2.6. Data presentation and statistical analysis

Data were analyzed using GraphPad Prism version 10.4.1 (GraphPad Software, San Diego, CA, USA) and figures were assembled in Affinity Designer 1.10 600bf (Serif Europe Ltd). Results are presented as mean ± standard deviation (SD), unless otherwise stated. Comparisons between two groups were assessed using a two-tailed unpaired *t*-test. For experiments involving multiple groups, one-way ANOVA followed by Bonferroni *post hoc* test was applied. For experiments involving multiple conditions or time points, two-way ANOVA with Bonferroni *post hoc* tests was used. The number of biological replicates (n) is indicated in each figure legend. Statistical significance was defined as *p* < 0.05 and significance values are given directly in the figures. Data collection and analysis were randomized, and investigators were blinded to group allocation during experiments and outcome assessment. Q-values for volcano plots were calculated by using the False Discovery Rate (FDR) with the original Benjamini-Hochberg method. The log2(fold change) cutoff was set to ± 2 (= 4-fold), and the −log10(q-value) threshold was set to 2 (= q-values lower than 0.01).

## 3. Results

### 3.1. Biochemical characterization and morphological assessment of murine *Peds1* and *Gnpat* KO eyes

As outlined in the introduction, the eyes are among the organs most severely affected by complete ether lipid deficiency. However, the specific impact of selective plasmalogen loss remains unexplored. To address this gap, we set out for a basic characterization of the eyes of *Peds1* KO mice to assess the role of plasmalogens and compared them to the eyes of WT littermates and *Gnpat* KO mice, which served as a previously described model of total ether lipid deficiency. Eye weights were determined and analyzed both as absolute values (**Fig. 1a**) and relative to total mouse body weight (**Fig. 1b**). Both parameters revealed a strong and significant reduction in *Gnpat* KO mice compared to *Peds1* KOs and, for absolute weight, also in the comparison with WT eyes. (**Fig. 1a**). Absolute eye weight in *Peds1* KO mice was decreased when compared to WT, albeit to a much lesser extent than in *Gnpat* KO. This reduction was absent when normalizing eye weight to body weight (**Fig. 1a, b**).

**Fig. 1:**
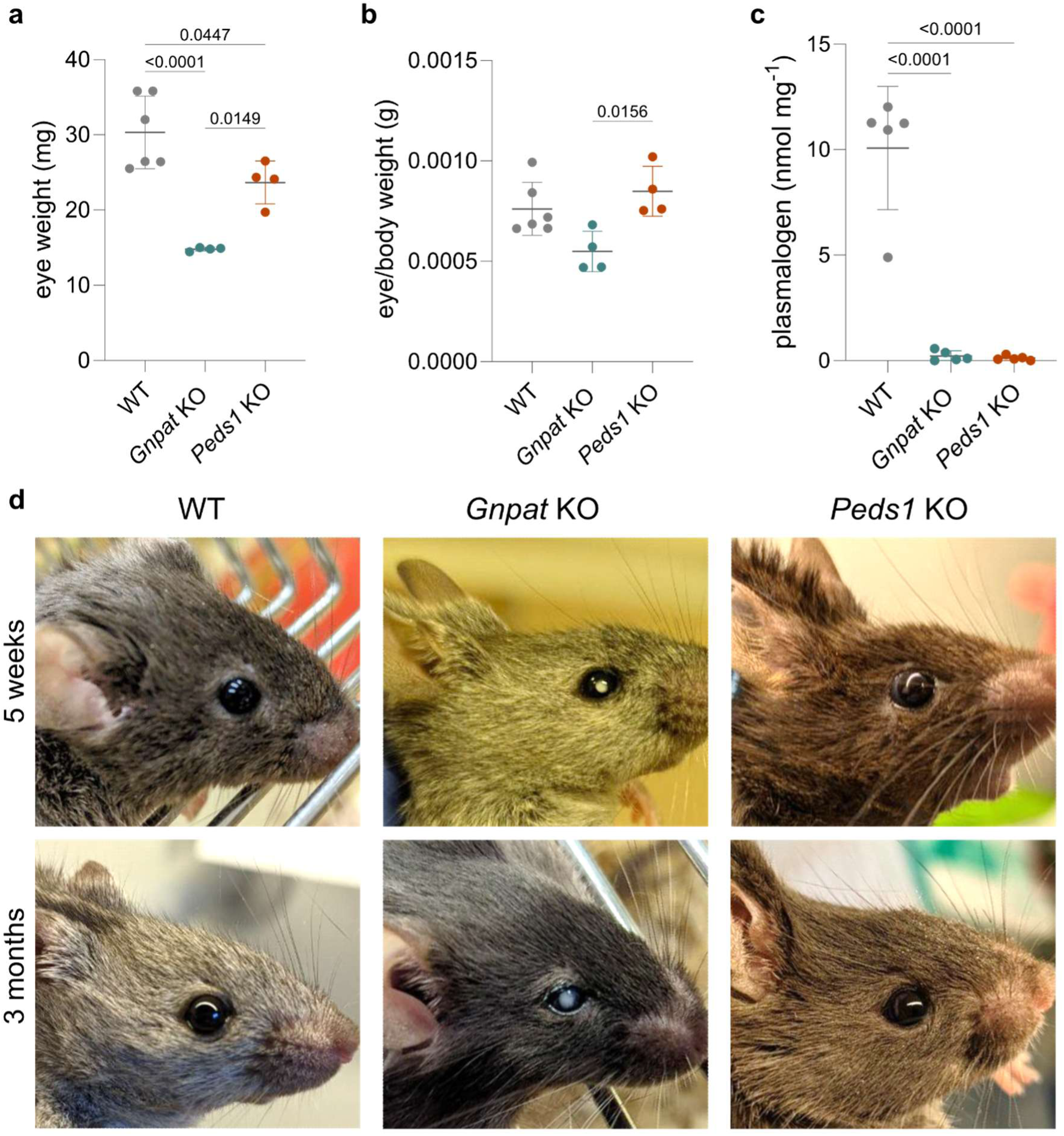
Weight analysis, biochemical phenotyping and cataract evaluation of murine Gnpat-and Peds1-deficient eyes. **a** Eye weights of WT, Gnpat KO and Peds1 KO mice at 3 months of age and **b** ratio of eye to body weight of the same animals (WT: n = 5, 2 females, 3 males; Gnpat KO: n = 4, 2 females, 2 males; Peds1 KO: n = 4, 2 females, 2 males). **c** Plasmalogen levels were quantified according to our established protocol using acid cleavage and derivatization to a fluorescent hydrazone [35] in the eyes of 6-month-old WT, Gnpat KO and Peds1 KO mice (n = 5). **a-c** Data are shown as mean ± SD, statistical analysis was performed by one-way ANOVA with Bonferroni post hoc test. **d** Representative images of eyes of WT, Gnpat KO and Peds1 KO at 5 weeks (upper panels) and 3 months of age (lower panels) are shown.

Next, we assessed ocular plasmalogen content in all three genotypes (**Fig. 1c**). WT eyes exhibited mean plasmalogen levels of 10.1 ± 2.9 nmol/mg tissue, whereas levels in both KO strains were significantly reduced approaching the detection limit (mean plasmalogen levels: *Gnpat* KO: 0.22 ± 0.25 nmol/mg; *Peds1* KO: 0.12 ± 0.11 nmol/mg, p < 0.0001, n = 5 per genotype). PEDS1 enzyme activity did not surpass detection limits in all analyzed eyes (limit of detection (LOD) = 0.2 pmol mg-1 min-1). This is in analogy to a previous study where out of 12 tested murine tissues only kidney, spleen, colon, and lung yielded detectable activity (ranging between 0.44 pmol mg^−1^ min^−1^ and 1.29 pmol mg^−1^ min^−1^) [35].

Basic phenotyping of *Peds1* KO mice by the International Mouse Phenotyping Consortium (IMPC) had identified an abnormal eye and retina morphology, as well as abnormal retinal pigmentation (https://www.mousephenotype.org/data/genes/MGI:2142624). Because cataracts, a fully penetrant feature of the human disease RCDP and several KO mouse models mimicking RCDP [23,24,42], were not listed among the observed *Peds1* KO phenotypes, we established a *Peds1* KO mice colony on a C57BL/6 x CD1 genetic background (n = 23, 11 females and 12 males), monitored cataract development up to six months of age, and compared the animals with age-matched *Gnpat* KO mice (n = 23, 8 females and 15 males) and littermate WT mice (n = 30, 14 females and 16 males). No cataracts were observed in *Peds1* KO mice at any time point including older animals, whereas all *Gnpat* KO mice exhibited them at every time point tested (**Fig. 1d**, representative pictures at 5 weeks and 3 months shown). Consistently, *Peds1* KO mice on pure C57BL/6N background also showed no cataracts, even up to 15 months.

### 3.2. Lipidomic analysis of the eyes of *Peds1* KO versus *Gnpat* KO and WT mice

Prompted by the absence of cataracts despite the complete lack of plasmalogens in *Peds1* KO eyes, we characterized their ocular lipid composition in more depth. In lipidomics measurements, we were able to obtain a total of 160 high-quality annotations for glycerophospholipids (139 species) and sphingolipids (21 species) (**Fig. 2a**). The former category was further stratified into 39 ether-linked glycerophospholipids and 100 ester-linked species (**Fig. 2a**). In WT eyes, PC constituted the most abundant subclass, followed by PE(P), PE and PS. Among ether lipids, a minor contribution of PE(O) and only traces of PC(O) and PS(O) were detected (**Supplementary Table 1**). PCA revealed distinct clustering of the genotypes with components 1 and 2 accounting for 40.9% and 31.3% of the total variance, respectively, driven most strongly by the lipid species PE(18:0/20:4), PE(P-18:1/18:1) and PE(O-18:1/18:1) (**Supplementary Fig. 1a**). Hierarchical cluster analysis of the top 50 features further confirmed the clear separation between *Gnpat* KO, *Peds1* KO, and WT groups (**Supplementary Fig. 1b**). *Gnpat* KO (**Fig. 2b**) and *Peds1* KO (**Fig. 2c**) did lead to strong and highly genotype-specific alterations of the molecular lipid species composition. Significant effects were observed mostly in phosphoethanolamine/-choline-linked subclasses (PE, PE(P), PE(O), PC(O)). The impact elicited by the two KOs was tightly localized to these subclasses and very pronounced, as indicated by the changes in subclass abundance (**Supplementary Table 1**) and the high number of altered individual species within each affected subclass (**Fig. 2b, c**). For ethanolamine-linked glycerophosphospecies, both KOs deviated strongly from the WT pattern: *Gnpat* KO eyes (**Fig. 2d**, green signals) completely lacked all annotated ethanolamine-containing ether lipids (PE(O) and PE(P)) compared to WT (**Fig. 2d**, gray signals) and as a compensatory mechanism had increased levels in phosphatidylethanolamines (PE), with 13 of the annotated 28 PEs being significantly upregulated. In contrast, *Peds1* KO eyes (**Fig. 2d**, red signals) had a PE distribution that was indistinguishable from WT. Here, we observed a selective depletion of PE(P) species, which was compensated by a concomitant upregulation of the corresponding plasmanyl lipid (PE(O)) species. This was observed for all 13 plasmanyl/plasmalogen pairs annotated in our analysis (e.g. PE(P-36:2) in WT and PE(O-36:2) in *Peds1* KO: high signal for both species among the 13 paired signals; PE(P-38:3) in WT and PE(O-38:3) in *Peds1* KO: low signal for both species among the 13 paired signals).

**Fig. 2:**
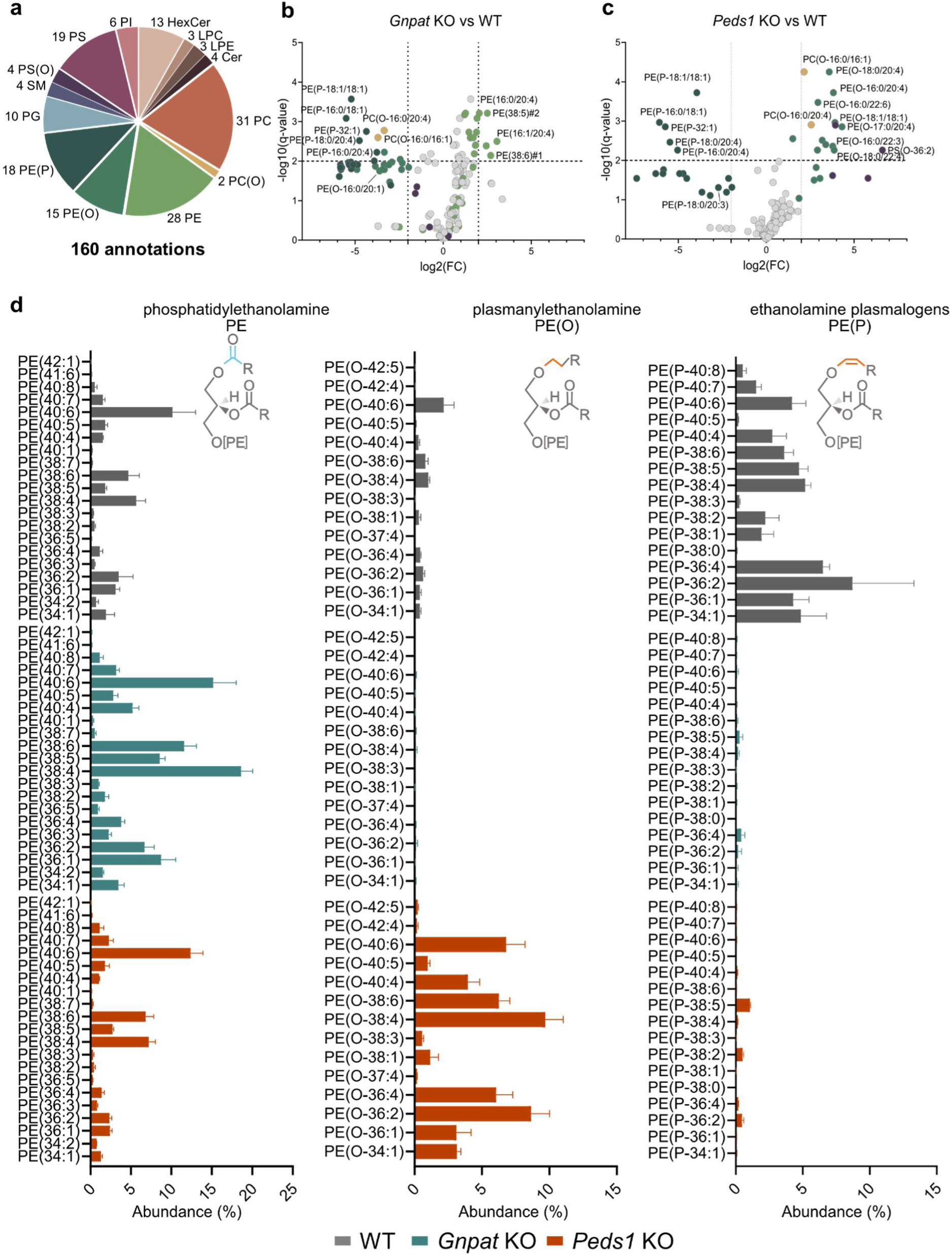
Lipidomic analysis of phospholipid subclasses in Peds1 KO, Gnpat KO and WT eyes. **a** 160 sphingo- and glycerophospholipid species corresponding to 14 lipid subclasses were annotated. Volcano plots of Gnpat KO **b** and Peds1 KO **c** versus WT eyes. For space reasons only selected lipid species are labeled. Lipid subclasses containing at least one significantly regulated species are shown in color according to **a**. Light gray dots denote lipids from subclasses of which no member was significantly altered. **d** Distribution of selected ethanolamine-based glycerophospholipids divided into phosphatidylethanolamines (PE, left), plasmanylethanolamines (PE(O), center) and plasmalogens (PE(P), right), in WT (gray, top), Gnpat KO (green, center) and Peds1 KO (red, bottom) eyes of 6-month-old male mice. Isobaric molecular species were condensed to their total carbon and double bond numbers (extended data can be found in the supplement). Relative abundances (%) were calculated by normalizing each species to the sum of all detected PE, PE(O), and PE(P) species. Data is presented as the mean ± SD. n = 5 per genotype, for all panels. Statistical analysis of volcano plots in **b** and **c** was performed by multiple unpaired t tests (original false discovery rate (FDR) method by Benjamini and Hochberg).

In conclusion, while the KO of *Gnpat* disrupts ether lipid biosynthesis and leads to their quantitative depletion that is accompanied by a compensatory upregulation of ester-linked lipids, the selective loss of plasmalogens in *Peds1* KO results in the accumulation of corresponding plasmanyl lipids with otherwise similar molecular composition.

### 3.3. Histological analysis of murine eyes deficient in all ether lipids versus selectively in plasmalogens

A previous report showed a marked ocular phenotype including microphthalmia in *Gnpat* KO mice [24] which is in line with our results (**Fig. 1a, b, d**). However, knowledge on the impact of selective plasmalogen deficiency on the eye has so far been limited to an IMPC basic screen where abnormal eye and retina morphology as well as disturbed retinal pigmentation were identified (https://www.mousephenotype.org/data/genes/MGI:2142624).

To directly compare the phenotypic consequences of total versus subclass-specific ether lipid deficiency in greater detail, we analyzed H&E-stained eye sections of male and female mice of both KO strains and compared them to WT control eyes (**Fig. 3a-f**). As previously described, the lens in *Gnpat* KO mice was significantly smaller and abnormally shaped compared to WT (**Fig. 3a, e**), while *Peds1* KO eyes appeared phenotypically indistinguishable from WT (**Fig. 3a, c**). In all three genotypes, the anterior portion of the lens was lined by a single-layered epithelium and a capsule. Both structures showed thickening in the area of the pupil in *Gnpat* KO mice (**Fig. 3e**, insert) but not in *Peds1* KO (**Fig. 3c**, insert) or WT animals (**Fig. 3a**, insert).

**Fig. 3:**
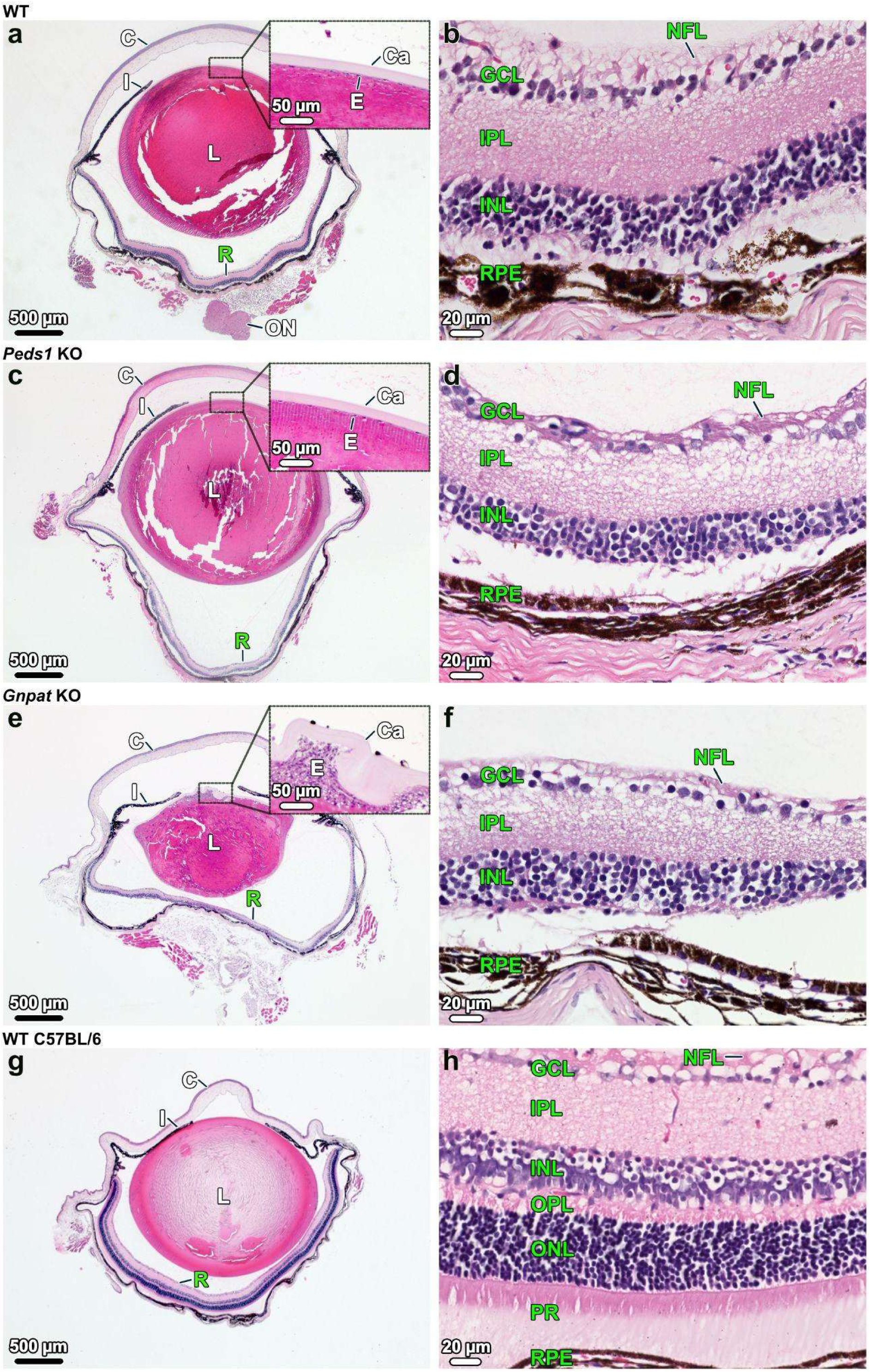
Histological analysis of Peds1-deficient eyes. Evaluation of eye histology in WT (**a**, **b**), Peds1 KO (**c**, **d**), Gnpat KO (**e**, **f**) mice (all on mixed C57BL/6 x CD1 genetic background), as well as in WT C57BL/6N mice (**g**, **h**). In WT and both KO models, the eyes consisted of the cornea (C), the iris (I), the lens (L), and the retina (R). **a, c, e** depict sagittal sections through the eyeball. The inserts show the anterior segment with the lens epithelium (E) and the lens capsule (Ca). **b, d, f** show the posterior segment, including the retina and its layers. All sections were H&E stained. ON = optic nerve; RPE = retinal pigmented epithelium, IPL = inner plexiform layer; INL = inner nuclear layer; GCL = ganglion cell layer; NFL = nerve fiber layer; OPL = outer plexiform layer; ONL = outer nuclear layer; PR = photoreceptors. One representative image of n = 1-4 is shown (WT: C57BL/6 x CD1, n = 4, 2 females, 2 males; WT C57BL/6N, n = 1 female; Peds1 KO: n = 3, 2 females, 1 male; Gnpat KO: n = 2, 1 female 1 male).

In particular, structural disruption and vacuolization of the lens epithelium were observed in the pupil area and beneath the iris of *Gnpat* KO mice - a feature that was not seen in the other two genotypes (**Fig. 4a, b** and for comparison inserts in **Fig. 3a, c**). No other differences in the structure of the cornea (C) and iris (I) were visible between WT and both KO strains (**Fig. 3a, c, e**). Serial sections showed that WT, *Peds1* KO and *Gnpat* KO mice all possessed an intact optic nerve (ON) (shown only for WT in **Fig. 3a**). In all three genotypes and both in 5 week- and 3-month-old mice, the retina showed impaired layering and consisted of a retinal pigment epithelium (RPE), an inner nuclear layer (INL), an inner plexiform layer (IPL), a ganglion cell layer (GCL), and a nerve fiber layer (NFL). However, the outer plexiform layer (OPL) and the outer nuclear layer (ONL) as well as photoreceptors (PR) were absent (compare **Fig. 3b, d, f** with the retina of a WT mouse on pure C57BL/6N background with normal retinal architecture (**Fig. 3g, h**)).

**Fig. 4:**
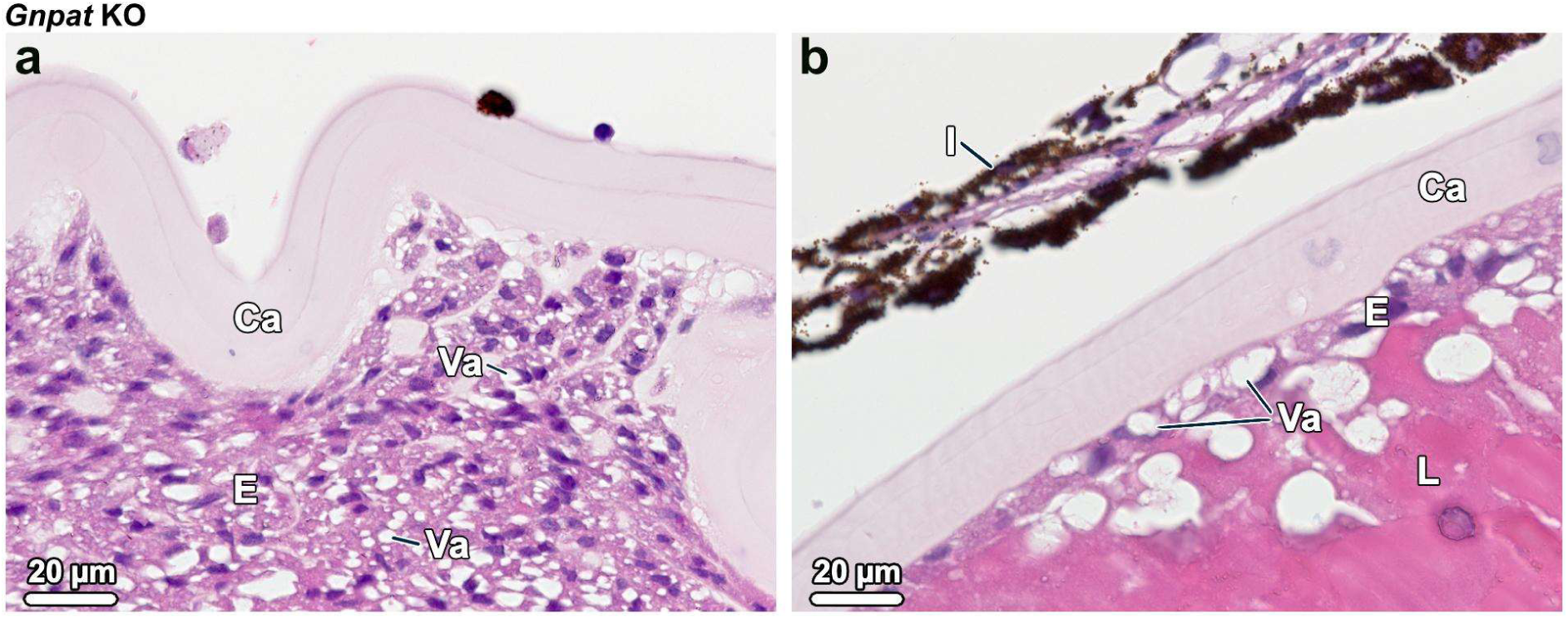
Histological examination of the Gnpat-deficient lens epithelium. Note that the lens epithelium (E) was thickened in the pupil area **a**, whereas it was single-layered below the iris (I) **b**. Ca = capsule, E = lens epithelium, I = iris, L = lens, Va = vacuoles. The sections were H&E stained. Representative image of a 3-month-old male mouse (n = 2).

Summarized, our analysis revealed striking lens differences elicited by the total ether lipid deficiency in *Gnpat* KO mice when compared to WT, while *Peds1* KO animals had normal lens structure. In addition, we found a disturbed retinal layer structuring in both KOs and the WT animals on mixed C57BL/6 x CD1 genetic background.

### 3.4. Assessment of the impact of selective plasmalogen deficiency on fertility and survival

*Gnpat* KO males have been reported to be completely infertile, whereas females exhibit significantly reduced fertility: only one out of seven females became pregnant after mating with WT males, and the two resulting pups died shortly after birth. Examination of genital structures revealed major changes in ovarian follicles, glassy membranes and corpora lutea [24]. A recent study meanwhile even suggested complete infertility for female *Gnpat* KO mice [28]. To gain additional insights into fertility and pup survival of the *Gnpat* KO mouse model, we examined 60 litters from *Gnpat* heterozygous (het) matings on the C57BL/6 x CD1 mixed genetic background (Fig. 5). In these litters a total of 453 animals were born with 378 pups surviving weaning (mortality rate until weaning: 16.6%). Mean litter size was 6.3 ± 2.2 pups, with a significant deviation of the number of KO pups from Mendelian genotype ratios (WT: 29 ± 20%, *Gnpat* het: 52 ± 26%, *Gnpat* KO: 17 ± 18%, WT versus KO: p = 0.0213). When stratifying the offspring by sex no significant deviation was found (WT: 56 females/55 males; *Gnpat* het: 93 females/108 males; *Gnpat* KO: 23 females/43 males; WT female to male: p = >0.999, *Gnpat* het female to male: p = >0.999, *Gnpat* KO female to male: p = >0.999). Preliminary data from the IMPC did not suggest an infertility or subfertility phenotype in the *Peds1*KO strain (https://www.mousephenotype.org/data/genes/MGI:2142624). To systematically assess reproductive parameters, we analyzed 15 litters from *Peds1* het breeding pairs on C57BL/6 x CD1 background (Fig. 5). In these litters a total of 103 animals were born and 90 of the pups survived weaning (mortality rate until weaning: 13%). The mean litter size was 6.0 ± 2.2 pups, with ratios indistinguishable from Mendelian ratios (WT: 24 ± 17%, het: 44 ± 24%, KO: 32 ± 18%, WT versus KO: p = 0.799), as well as a balanced sex distribution (WT: 10 females/13 males; *Peds1* het: 22 females/18 males; *Peds1* KO: 11 females/16 males; WT female to male: p = >0.999, *Peds1* het female to male: p = >0.999, *Peds1* KO female to male: p = >0.999). For the *Peds1* strain, we also analyzed 36 litters from het matings on pure C57BL/6N background (Fig. 5). In these litters a total of 275 animals were born and 236 of the pups survived weaning yielding a mortality rate until weaning of 14.2% and a mean litter size of 7.6 ± 2.3 pups. No significant deviation from Mendelian ratios was observed, despite a trend to reduction in *Peds1* KO pups (WT: 28 ± 23%, het: 56 ± 24%, KO: 16 ± 13%, WT versus KO: p = 0.563) and unperturbed sex ratios (WT: 35 females/32 males; *Peds1* het: 69 females/58 males; *Peds1* KO: 20 females/22 males; WT female to male: p = 0.989, *Peds1* het female to male: p = >0.999, *Peds1* KO female to male: p = >0.999). To further assess the fertility of plasmalogen-devoid animals on pure C57BL/6N background, we bred *Peds1* KO males with *Peds1* KO females. These matings indeed yielded offspring; breeding was discontinued after five litters had been obtained (28 offspring, of which two did not survive weaning (mortality rate until weaning: 7.1%)). The mean litter size was 5.6 ± 1.4 and did not differ significantly from that of heterozygous breedings (p = 0.067). There was a non-significant male bias (9 females/17 males). In general, the sex distribution did not differ significantly from that obtained from het x het breedings (p = 0.1103).

**Fig. 5:**
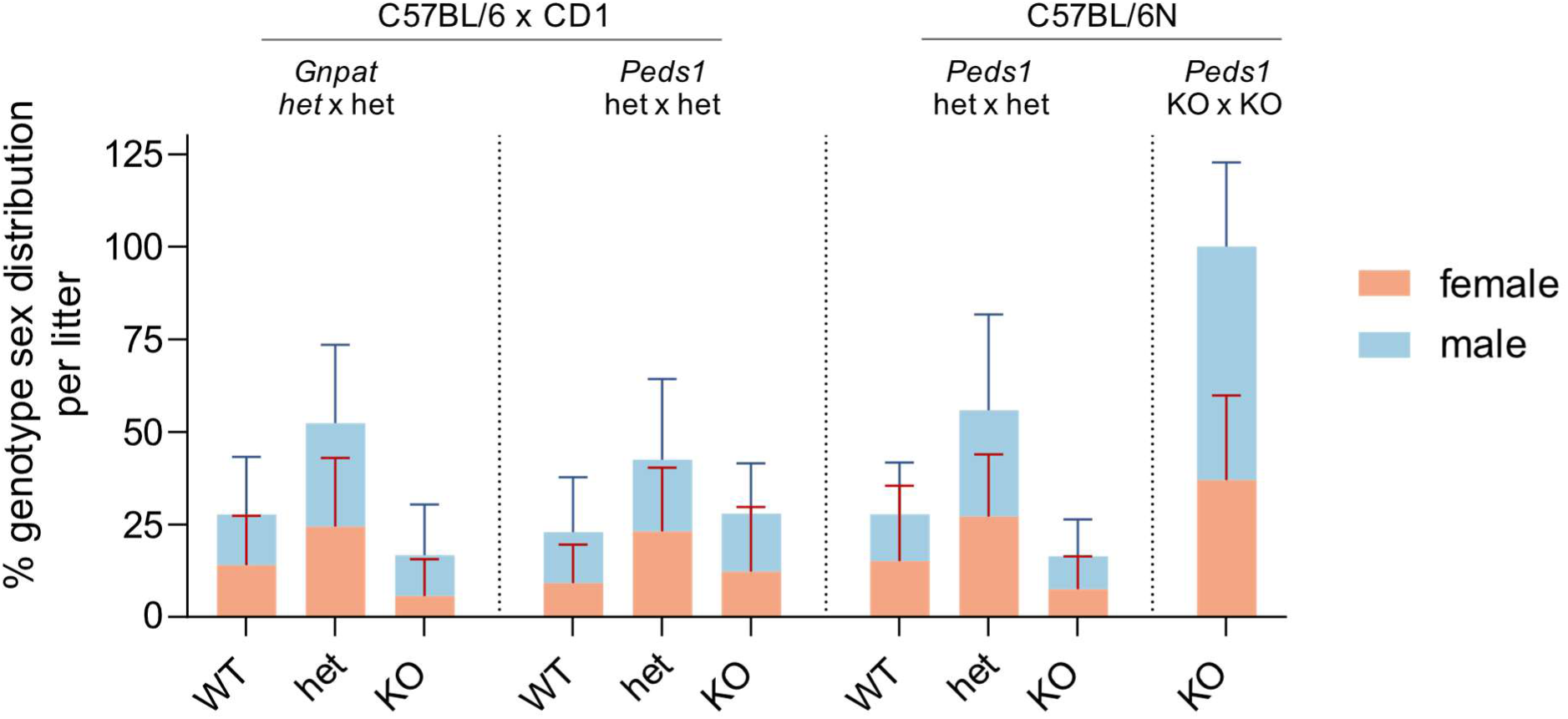
Analysis of litters obtained from mating (from left to right) of heterozygous Gnpat animals (n = 60 litters) and Peds1 animals (n = 15 litters) on mixed C57BL/6 x CD1 background, as well as of heterozygous Peds1 (n = 36 litters) and homozygous Peds1 (n = 5 litters) on pure C57BL/6N background. Offspring are color-coded according to their sex, with orange indicating female pups and light blue representing male pups. het = heterozygous. Data are shown as mean ± SD.

In contrast to the profound reproductive defects in *Gnpat* KO mice, *Peds1* KO animals displayed normal fertility, Mendelian distribution and pup survival across both mixed and pure backgrounds. Moreover, *Peds1* KO mice produced viable litters with parameters indistinguishable from het matings.

## 4. Discussion

This detailed molecular and phenotypic comparison of mice totally deficient in ether lipids (*Gnpat* KO) and mice deficient in plasmalogens only (*Peds1* KO) shows for the first time that plasmanyl lipids retained in *Peds1* KO are sufficient to alleviate the pronounced eye and fertility defects characteristic of *Gnpat* KO mice. Eyes and specifically the lenses of *Peds1* KO mice were found indistinguishable from WT in overall visual assessment and histology despite significant differences in their phospholipidome. Furthermore, *Peds1* KO mice were able to maintain physiological functioning of the reproductive system in both males and females, thus negating an essential role of plasmalogens in the involved processes ranging from spermatogenesis and oogenesis to implantation, gestation and parturition.

Different mouse models mimicking RCDP, which is characterized by a total deficiency in all ether lipid subclasses, have been available for decades and were extensively characterized [24,25,43–46]. In murine *Gnpat* KO models specifically, the eye phenotype involves multiple structural abnormalities. Besides the distinct dense bilateral cataracts, microphthalmia and dysgenesis of the anterior segment was found. Furthermore, morphological changes were observed in lens epithelial cells, together with swelling and disorganization of lens fiber cells. Alterations in the thickness of both the anterior and posterior portions of the lens capsule have also been reported. Additional defects comprise the formation of lenticonus abnormally adhering to the corneal endothelium [22,24]. Retinal alterations have also been described, in particular an increased thickness of Bruch’s membrane accompanied by structural and functional abnormalities of the RPE. These include vacuolization, regions of both hypo- and hyperplasia, altered pigmentation, and the accumulation of photoreceptor-derived degradation products [22,24]. Here, we could reproduce bilateral cataracts, microphthalmia, thickening of the lens capsule, as well as disorganization and vacuolization of the single-layered cubic lens epithelium in the anterior segment of the eye (**Figs. 1, 3**, and **4**). Other features, such as vacuolization of the RPE and changes in optic nerve myelination described in [24], were not observed in our model system. Unlike ref. [24], which had used osmium to fix lipid-rich structures such as myelin, we used paraformaldehyde, and these preparation differences presumably limited our ability to evaluate the extent of optic nerve myelination.

*Peds1* KO mice had been made available as *Tmem189tm1a(KOMP)Wtsi* mice by the EMMA consortium and have since transformed into an essential tool to study ether lipid biology. We utilized them to set up reliable mass spectrometric protocols for the unequivocal annotation of ether lipids in lipidomic analyses [reviewed in 18,40] and to investigate the impact of the lack of plasmalogens on the murine organism [16,34]. In addition, we could recently show profound changes in the haematological and immune system of these mice [see preprint 19]. Very recently, the first human bi-allelic *PEDS1* variant (chr20: g.50153533AG > A(hg38); NM_199129.4: c.104delC;p.Ala35Valfs*16) was described in two male patients suffering from microcephaly, developmental delays and, in one case, also from congenital cataracts [7]. This finding was particularly striking, as cataracts are one of the most penetrant symptoms of RCDP in humans. The only available baseline ocular phenotype annotation in the corresponding *Peds1* KO model performed by the IMPC listed changes in eye and retina morphology as well as disturbed retinal pigmentation but did not include cataracts (https://www.mousephenotype.org/data/genes/MGI:2142624). Prompted by this discrepancy, we set out to perform an in-depth characterization of *Peds1*-deficient eyes and now show that, in contrast to the strong alterations observed in *Gnpat* KO eyes, *Peds1* KO mice did not develop cataracts in any cohort examined (**Fig. 1**). Specifically, in both a mixed genetic background and a pure C57BL/6N cohort, we never observed any cataracts up to 15 months of age and detected no changes in the lens epithelium or lens capsule (**Fig. 3**).

Mass spectrometry-based lipidomics analysis of ocular phospholipids revealed that, among 160 annotated lipid species belonging to the two lipid categories of glycerophospholipids and sphingolipids, KO-elicited alterations were strictly confined to glycerophospholipids, affecting the subclasses of PE, PE(O), PE(P), PC(O) and PS(O). Under WT steady-state conditions high amounts of PE plasmenyl lipids (PE(P), 51.2% of total PE) compared to plasmanyl lipids (PE(O), 6.9% of total PE) were observed. This is in contrast to human lenses which contain high amounts of plasmanyl lipids [47]. The same study also identified PS(O) species in human lenses, aligning with ref. [14], which specifically reports PS(O-16:0/18:1) and PS(O-18:1/18:1). Our study detected four PS(O) species of which two likely represent the same molecular entity (PS(O-16:0/18:1) and PS(O-36:2)). Further prominent species in our murine study were PE(O-18:1/18:1), already identified to be most abundant in human lenses [14], and PE(P-18:1/18:1), previously found to be the second most abundant glycerophospholipid in mouse lenses [14]. Both of these lipids contributed strongly to the separation of WT, *Peds1* KO and *Gnpat* KO mice in the PCA. In addition, ether lipid species containing arachidonic side chains at *sn*-2, that had been described to be major species in mouse retinal epithelium [10], appeared prominently in our study (PE(P-16:0/20:4), PE(P-18:0/20:4)). In order to investigate whether the high plasmenyl/plasmanyl ratio and the low PEDS1 activity are a consequence of low expression levels of *Peds1*, we analyzed available retinal gene expression data [48] for *Peds1*, *Gnpat* and five additional genes crucially involved in ether lipid biosynthesis (**Fig. 6**). It is interesting to note that, despite the relatively low expression level of *Peds1* and its very low enzymatic activity, the conversion from plasmanyl to plasmenyl species appears to be highly efficient in this ether lipid-enriched tissue.

**Fig. 6:**
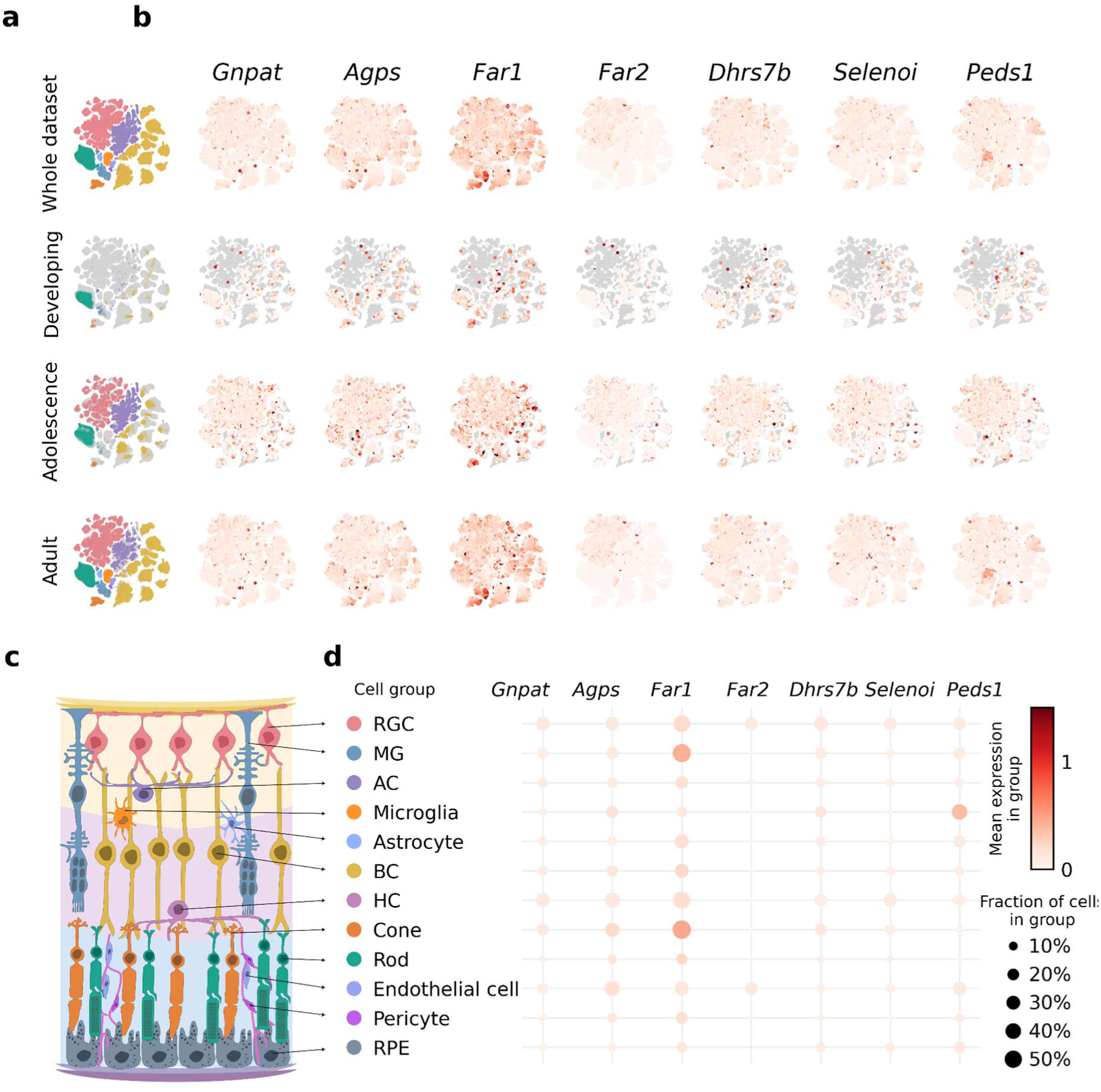
Transcriptomics analysis of selected genes associated with ether lipid metabolism in mouse retina. **a** Cell groups present in the mouse retina single-cell atlas during mouse development [48]. The high dimensional transcriptomics data is visualized as a Uniform Manifold Approximation and Projection (UMAP), retaining the overall topology of the data. Mouse development was binned into developing (up until age P14), adolescence (up until P28), and adult (after P28). Samples are colored based on their cell group annotation, or in gray if they are not part of the age bin. **b** Expression of selected ether lipid metabolism-associated genes during mouse development projected into UMAP space as 2D histogram showing their mean expression in each hexagonal bin. **c** Topology and cell types in mouse retina. RGC = retinal ganglion cells; AC = amacrine cells; BC = bipolar cells; HC = horizontal cells; MG = Müller Glia; RPE = retinal pigment epithelium. **d** Mean expression of selected ether lipid metabolism-associated genes across cell types of the mouse retina over all developmental stages.

As expected, *Gnpat*-deficient eyes lacked all ether-linked species. In their place, PE species were strongly upregulated to 97.7% of total PE but differed in side-chain composition and unsaturation pattern from the composition of the physiological plasmalogen pool in WT eyes. Such a compensatory remodeling is consistent with previous findings where, despite loss of ether lipid metabolic enzymes, total ethanolamine glycerophospholipid levels were kept constant [49,50] most probably to maintain membrane properties.

In contrast, *Peds1* KO eyes preserved WT levels of ester lipids and instead compensated for their selective loss of plasmalogens by increased levels of plasmanyl lipids (50.3% of total PE) with side-chain distribution and desaturation pattern strongly recapitulating that of the absent plasmalogens (**Fig. 2d**). This goes hand in hand with previous findings [16] and indicates that PEDS1 accepts a wide range of PE(O) species as substrates for desaturation to PE(P) and that consequently, in the absence of the enzyme, these precursors accumulate. Other accumulated species in *Peds1* KO mice were two of the four detected plasmanylserine species (PS(O-36:2) and PS(O-40:6), **Fig. 2c**). Together, this shows that while plasmalogens are the predominant phosphoethanolamine lipid subclass in the WT eye, when they cannot be synthesized, plasmanyl lipids are sufficient to maintain normal eye development and structure. A defined role of ether lipids in the lens has never been assigned, but their involvement in the maintenance of lens transparency and growth has been hypothesized [22]. This is supported by the finding of *gnpat* expression in *X. laevis* proliferating lens forming cells [51]. These cells eliminate their nucleus in order to allow for the lens to be “organelle-free” and thereby transparent. PE(P) were described as main components in young fibers in lens membranes, hinting at a role of ether lipids and their biosynthetic enzymes like GNPAT in lens fiber differentiation and elongation. Older more central fibers were shown to rely on considerable amounts of sphingomyelins, which increase with age and also in the cataractous lens while plasmalogen levels decline under these conditions [51].

A striking and unexpected finding in all analyzed eyes, including WT, from animals bred on the mixed background (C57BL/6 x CD1) was the outright absence of the outer plexiform layer (OPL), the outer nuclear layer (ONL) and the photoreceptor layer (**Fig. 3**), indicating that this phenotype was background-dependent rather than genotype-specific. As these three layers of the retina are essential for vision, our results suggest that mice on the mixed background are functionally blind, as no image-forming light signals can be transmitted to the brain. Previous analyses of *Gnpat* KO eyes (on a pure C57BL/6 background) did not show this reduction of the retinal layers [24]. We also tested C57BL/6N WT mice and found the retinal layer structure to be intact in this setting (**Fig. 3h**). A similar phenotype involving visual impairment and thinning of the outer nuclear layer of the retina was identified in the surfactant protein genes *Sftpa1* and *Sftpd* KO mice after outbreeding with Black Swiss mice carrying a mutation in phosphodiesterase-6 (PDE6). Backcrossing of those lines onto a pure C57BL/6 background restored the physiological structure of the retina as well as vision [52]. Our mixed background contains parts of CD1 mice, which are related to Black Swiss mice, as this enabled survival of KO animals. Backcrossing to a pure C57BL/6 background is therefore not feasible for the *Gnpat* KO strain in our study. Collectively, these observations support the presence of a segregating retinal degeneration allele from the CD1/Black Swiss lineage as the most plausible cause of the retinal phenotype on the mixed background.

*Gnpat* male KO animals are infertile, while females are described as subfertile or infertile [24,27,28]. In males, testes show defective spermatogenesis, characterized by severe germ-cell loss, seminiferous tubule atrophy, and a paucity of elongating/elongated spermatids and mature sperm, associated with disrupted blood-testis barrier integrity and cytoskeletal disorganization [24,27]. In females, ovarian alterations, including abnormal follicle morphology, impaired ovulation and reduced fertility have been reported [24,28]. For *Peds1* KO mice on pure C57BL/6N background no overt reproductive system phenotype had been listed by the IMPC (https://www.mousephenotype.org/data/genes/MGI:2142624), which is consistent with our findings after heterozygous *Peds1* matings of normal litter size (7.6 pups), moderate pup mortality (14.2%, between birth and weaning) and Mendelian genotype distribution. A numerical underrepresentation of KO pups at birth (16%) was observed, but normalized (32%) on the C57BL/6 x CD1 background (**Fig. 5**) where mortality of 13% was similar to the values obtained on the pure background. In *Gnpat* and *Agps* het matings on pure C57BL/6 background, severely reduced or even no KO mice had been found by other authors [53,54]. Outbreeding of the background to CD1 allowed us to obtain *Gnpat* KO animals from het matings in 17% of all weaned mice. Mortality rates between birth and weaning were comparable to *Peds1* breedings. The more severe outcome than that for *Peds1* KO mice on mixed background (**Fig. 5**) indicates a heavier impairment in total ether lipid deficiency than in selective plasmalogen loss. This is further evidenced by the fact that we obtained viable KO pups from *Peds1* KO x KO matings (on C57BL/6N background) with litter sizes of 5.6 ± 1.4 and lethality of only 7.1%. As seminolipid, a testis-specific sulfogalactosylglycerolipid essential for spermatogenesis and male fertility contains an alkyl chain at *sn*-1 [55] and can still be produced in the *Peds1* KO mouse but is naturally unavailable in the *Gnpat* KO mouse, this is in line with impaired male fertility in *Gnpat* KO animals (and other models of total ether lipid deficiency) while *Peds1* KO males are capable of reproduction. In females, the underlying mechanisms remain to be investigated in more detail.

In conclusion, our data show that in *Peds1* KO mice, retention of plasmanyl lipids is sufficient to preserve eye morphology and lens structure, as well as fertility and pup survival. In contrast, *Gnpat* KO mice with complete ether lipid deficiency exhibit severe ocular pathology and exhibit subfertility or infertility with markedly diminished KO pup survival. Future studies should define distinctive membrane properties of ether lipids and determine how these features, present in plasmanyl/plasmalogen lipids but not in diacyl ester analogues, support normal function in the eye and the reproductive system.

## Abbreviations

ACN: acetonitrile
AGPS: alkylglycerone phosphate synthase C cornea
Ca: lens capsule
DDA: data-dependent MS/MS acquisition
E: lens epithelium
ER: endoplasmic reticulum
FAR1/FAR2: fatty acyl-CoA reductase ½
GCL: ganglion cell layer
GNPAT: glyceronephosphate *O*-acyltransferase
het: heterozygous
HPC: high precision calibration
H&E: hematoxylin and eosin
I: iris
IMPC: International Mouse Phenotyping Consortium
INL: inner nuclear layer
IPA: isopropanol
IPL: inner plexiform layer
KO: knockout
L: lens
LC-MS: liquid chromatography-mass spectrometry
LOD: limit of detection
NFL: nerve fiber layer
ON: optic nerve
ONL: outer nuclear layer
OPL: outer plexiform layer
PAF: platelet-activating factor
PBS: phosphate-buffered saline
PCA: principal component analysis
PC(P): choline plasmalogen
PE: phosphatidylethanolamine
PEDS1: plasmanylethanolamine desaturase
PE(P): ethanolamine plasmalogen
PEX: peroxin
PFA: paraformaldehyde
PFCRD: peroxisomal fatty acyl-CoA reductase 1 disorder
PR: photoreceptors
PS: phosphatidylserine
R: retina
RCDP: Rhizomelic Chondrodysplasia Punctata
RPE: retinal pigment epithelium
SPF: specific pathogen free
SD: standard deviation
VA: vacuoles
WT: wildtype

## Acknowledgements

We are indebted to Nina Madl, Nikolas Andresen, Bianca Hafner, Christof Seifarth and Annabella Knab for expert technical assistance. We also thank the Wellcome Trust Sanger Institute Mouse Genetics Project (Sanger MPG) and its funders for providing the mutant mouse line *Tmem189tm1a(KOMP)Wtsi* and INFRAFRONTIER/EMMA (https://www.infrafrontier.eu/).

## Data availability

LC-MS/MS data is in the process of being deposited in a publicly accessible repository and will be available there prior to acceptance of the article. Other data will be made available upon reasonable request.

## CRediT author contributions

Conceptualization: FD, GG, JB, KW

Data Curation: ID, VJ, MJB, MAK

Formal Analysis: ID, VJ, MJB

Funding Acquisition: ID, MAK, KW

Investigation: ID, VJ, MJB, DK, JK, FD

Methodology: MJB, MAK

Project Administration: KW

Resources: MAK

Software: VJ, JK, MAK

Supervision: GG, JB, MAK, KW

Validation: ID, KW

Visualization: ID, MJB, JK

Writing – Original Draft: ID, MJB, KW

Writing – Review & Editing: ID, VJ, MJB, DK, JK, GG, FD, JB, MAK, KW

