## supplementary table and figure for "Beyond the surface: plasmalogens are dispensable for retinal integrity and fertility in the mouse"

### Abstract

Ether lipids and their subclass, the plasmalogens, are critical regulators of membrane organization, signaling, and stress responses in multiple tissues. Inborn errors in their anabolism cause severe multi-organ diseases such as Rhizomelic Chondrodysplasia Punctata and related peroxisomal disorders. The *Gnpat* knockout mouse model, characterized by total ether lipid deficiency, recapitulates key features of this disorder, including dense bilateral cataracts, microphthalmia, and infertility, but the specific contribution of different subclasses like plasmalogens remains elusive. The recent identification of the *Peds1* gene allows dissecting the impact of selective plasmalogen deficiency with retention of plasmanyl lipids, another ether lipid subgroup. Here, we performed the first side-by-side comparison of *Gnpat* and *Peds1* knockout in mice on a matched genetic background (C57BL/6 x CD1). In contrast to the situation in *Gnpat* knockout mice, plasmanyl lipids in *Peds1* knockout mice were sufficient to prevent cataract formation and maintain normal ocular structures, despite marked shifts in the ocular phospholipidome. Also, fertility and reproductive function were found to be preserved in *Peds1* knockout mice. Our data demonstrate that plasmanyl lipids can partially protect against the severe phenotypes observed in mouse models of total ether lipid deficiency; notably, the ocular and reproductive phenotypes were plasmalogen-independent, indicating that loss of the vinyl ether double bond is not the key determinant of all symptoms in human and murine ether lipid deficiency and can at least partly be compensated by plasmanyl lipids.

***Supplementary Table 1:*** *Mean relative abundance of lipid subclasses across WT,* Gnpat *KO and* Peds1 *KO eyes as determined by phospholipidomic analyses. Data are presented as mean percentage ± SD for n = 5 biological replicates per genotype. For each biological sample, the sum of peak intensities for all species within a specific lipid class was calculated and these class sums were then expressed as a percentage of the total integrated lipidome intensity (100%) within each individual replicate. Cer = ceramide; HexCer = hexosylceramide; LPC = lysophosphatidylcholine; LPE = lysophosphatidylethanolamine; PC = phosphatidylcholine; PG = phosphatidylglycerol; PI = phosphatidylinositol; PS = phosphatidylserin; SM = sphingomyelin.*

| **Lipid subclass** | **WT** (mean % ± SD) | ***Gnpat* KO** (mean % ± SD) | ***Peds1* KO** (mean % ± SD) |
|---|---|---|---|
| **Cer** | 3.35 ± 0.54 | 4.48 ± 0.39 | 3.91 ± 0.36 |
| **HexCer** | 11.19 ± 1.88 | 10.88 ± 1.28 | 10.81 ± 1.75 |
| **LPC** | 0.29 ± 0.02 | 0.56 ± 0.20 | 0.32 ± 0.01 |
| **LPE** | 0.12 ± 0.06 | 0.75 ± 0.58 | 0.11 ± 0.02 |
| **PC** | 22.12 ± 3.36 | 26.09 ± 1.36 | 22.34 ± 0.30 |
| **PC(O)** | 0.07 ± 0.03 | 0.00 ± 0.00 | 0.31 ± 0.12 |
| **PE** | 15.59 ± 2.62 | 32.54 ± 2.51 | 15.73 ± 0.59 |
| **PE(O)** | 2.51 ± 0.54 | 0.19 ± 0.13 | 17.45 ± 1.85 |
| **PE(P)** | 19.43 ± 2.12 | 0.53 ± 0.52 | 1.01 ± 0.14 |
| **PG** | 1.64 ± 0.50 | 2.57 ± 0.14 | 2.38 ± 0.39 |
| **PI** | 8.55 ± 1.59 | 8.88 ± 0.66 | 9.95 ± 0.78 |
| **PS** | 12.24 ± 1.12 | 9.38 ± 1.06 | 11.82 ± 0.66 |
| **PS(O)** | 0.04 ± 0.01 | 0.01 ± 0.01 | 0.87 ± 0.19 |
| **SM** | 2.87 ± 1.58 | 3.13 ± 0.79 | 3.00 ± 1.40 |
| **Sum** | 100.00 | 100.00 | 100.00 |

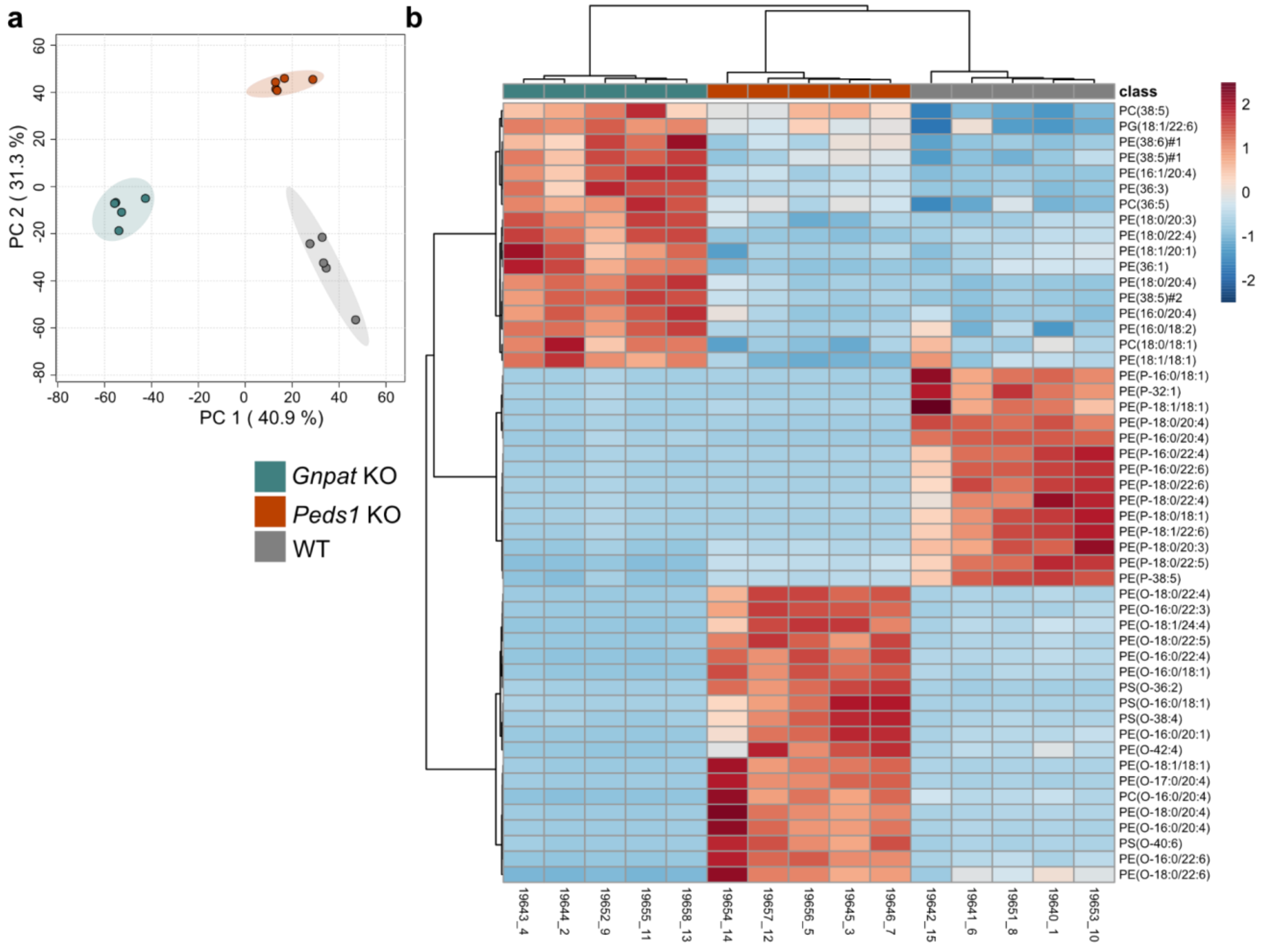


***Supplementary Fig. 1:*** *Multivariate statistical analysis of lipidomic analysis of selected lipid categories in WT,* Gnpat *KO and* Peds1 *KO eyes.* ***a*** *PCA score plot showing the separation of genotypes based on lipidomic profiles.* ***b*** *Heatmap and hierarchical cluster analysis of the top 50 lipid features. Each row represents an individual lipid species and columns represent individual biological replicates. The color scale indicates the normalized relative abundance of each lipid (red: high, blue: low, see scale legend). Samples were obtained from 6-month-old male mice (n = 5 per genotype for both panels). Statistical analysis and data visualization was performed using MetaboAnalyst 6.0. Data was sum normalized and autoscaled prior to the analysis. Distance was measured with Euclidean and clustering was performed with Ward method.*
